# Increased axonal bouton stability during learning in the mouse model of MECP2 duplication syndrome

**DOI:** 10.1101/186239

**Authors:** Ryan T. Ash, Paul G. Fahey, Jiyoung Park, Huda Y. Zoghbi, Stelios M. Smirnakis

## Abstract

*MECP2*-duplication syndrome is an X-linked form of syndromic autism caused by genomic duplication of the region encoding Methyl-CpG-binding protein 2. Mice overexpressing *MECP2* demonstrate altered patterns of learning and memory, including enhanced motor learning. Previous work associated this enhanced motor learning to abnormally increased stability of dendritic spine clusters formed in the apical tuft of corticospinal, area M1, neurons during rotarod training. In the current study, we measure the structural plasticity of axonal boutons in Layer 5 (L5) pyramidal neuron projections to layer 1 of area M1 during motor learning. In wild-type mice we find that during rotarod training, bouton formation rate changes minimally, if at all, while bouton elimination rate doubles. Notably, the observed upregulation in bouton elimination with learning is absent in *MECP2*-duplication mice. This result provides further evidence of imbalance between structural stability and plasticity in this form of syndromic autism. Furthermore, the observation that axonal bouton elimination doubles with motor learning in wild-type animals contrasts with the increase of dendritic spine consolidation observed in corticospinal neurons at the same layer. This dissociation suggests that different area M1 microcircuits may manifest different patterns of structural synaptic plasticity during motor learning.

**SIGNIFICANCE STATEMENT:** Abnormal balance between synaptic stability and plasticity is a feature of several autism spectrum disorders, often corroborated by in vivo studies of dendritic spine turnover. Here we provide the first evidence that abnormally increased stability of axonal boutons, the presynaptic component of excitatory synapses, occurs during motor learning in the MECP2 duplication syndrome mouse model of autism. In contrast, in normal controls, axonal bouton elimination in L5 pyramidal neuron projections to layer 1 of area M1 doubles with motor learning. The fact that axonal projection boutons get eliminated, while corticospinal dendritic spines get consolidated with motor learning in layer 1 of area M1, suggests that structural plasticity manifestations differ across different M1 microcircuits.

## INTRODUCTION

The rewiring of synaptic connections in neural microcircuits provides a compelling mechanism for learning and memory throughout development and adult life(Chklovskii et al., 2004). Two-photon imaging of fluorescently-labeled neurons has recently enabled the direct measurement of synaptic rewiring in vivo, revealing that new synapses form in motor cortex (M1) during motor training, and that the stability of these synapses correlates with how well the animal learns to perform the motor task (Xu et al., 2009; Yang et al., 2009). The layer 1 (L1) apical tuft dendritic spines that turn over during learning receive inputs from a range of sources, including L2/3, L5, and L6 cortical pyramidal neurons, thalamocortical neurons, and others. It is currently not known how synaptic inputs from axonal projections to area M1 behave during learning.

Experimental LTP and LTD paradigms in vitro can induce axonal bouton formation and elimination (Antonova et al., 2001; Becker et al., 2008; Bourne et al., 2013). In vivo, axonal boutons are spontaneously formed and eliminated in adult sensory cortex (De Paola et al., 2006; Majewska et al., 2006; Stettler et al., 2006; Grillo et al., 2013), while learning has been shown to alter bouton turnover in parallel fiber inputs to the cerebellum (Carrillo et al., 2013)and in orbitofrontal inputs to the medial prefrontal cortex (Johnson et al., 2016). However, bouton plasticity has yet to be measured in area M1 during motor learning to our knowledge. In this work we examine the turnover of boutons, the pre-synaptic component of synapses, in L5 pyramidal neuron axons that project to layer 1 of area M1.

Furthermore, we begin to assess whether learning-associated plasticity in inputs to area M1 is altered in the *MECP2* duplication model of autism. *MECP2* duplication syndrome is caused by a genomic duplication that spans the methyl-CpG-binding protein 2 (*MECP2*) gene and leads to a progressive X-linked disorder of intellectual disability, autism, spasticity, and epilepsy (Ramocki et al., 2010). Overexpression of the *MECP2* gene in mice produces a similar progressive neurological phenotype including autistic features (abnormal social behavior, anxiety, and stereotypies), spasticity, and epilepsy (Collins et al., 2004). Interestingly, before 24 weeks of age *MECP2*-duplication mice (Tg1) demonstrate a striking enhancement in motor learning and memory on the rotarod task (Collins et al., 2004). Previous work associated this enhanced learning with an increase in the formation and stabilization of dendritic spine clusters in apical dendritic tufts of corticospinal neurons in primary motor cortex (M1) (Ash et al., 2017), pointing to a possible mechanism for altered learning and memory in these animals.

MeCP2 and other autism-associated proteins contribute to the development of mature axons and presynaptic structures (Antar et al., 2006; Belichenko et al., 2009; Degano et al., 2009; Chen et al., 2014; Garcia-Junco-Clemente and Golshani, 2014). Presynaptic electrophysiological function has been shown to be altered in *MECP2*-duplication mice (increased paired pulse facilitation, Collins et al., 2004) and other autism mouse models (Deng et al., 2013), and mice with mutations in the proteins mediating presynaptic plasticity often demonstrate autistic features (Blundell et al., 2010). Long term depression (LTD), a form of synaptic weakening that has a major pre-synaptic component (Collingridge et al., 2010), has been shown to be defective in several models of autism (D’Antoni et al., 2014). These findings implicate pre-synaptic dysfunction in autism, but axonal bouton structural plasticity has not been explored directly in a model of autism to our knowledge.

We measured learning-associated axonal bouton structural plasticity in layer 1 of mouse M1 during rotarod training in the Tg1 mouse model of the *MECP2* duplication syndrome and compared with wild-type (WT) littermates. We found that the rate of bouton formation does not change significantly with rotarod training in either genotype, remaining approximately the same as the spontaneous bouton formation rate at rest. In contrast, bouton elimination rate is dramatically accelerated during rotarod learning in WT mice, whereas this effect is completely abolished in *MECP2*-duplication mice. This supports the argument that increased synaptic stability manifests in the MECP2-duplication syndrome during learning (Ash et al., 2017).

## MATERIALS & METHODS

### Animals

FVB-background *MECP2*-duplication (Tg1) mice (Collins et al., 2004), were crossed to C57 thy1-GFP-M (Feng et al., 2000) homozygotes obtained from Jackson Laboratories, to generate male F1C57;FVB *MECP2*-duplication;thy1-GFP-M mice and thy1-GFP-M littermate controls.

### In vivo two-photon imaging

All surgeries and imaging were performed blind to genotype. At least two weeks prior to the first imaging session (~12-14 week-old-mice), a 3 mm-wide opening was drilled over motor cortex, centered at 1.6 mm lateral to bregma (Tennant et al., 2011), and a glass coverslip was placed over the exposed brain surface to allow chronic imaging of neuronal morphology (Mostany and Portera-Cailliau, 2008; Holtmaat et al., 2009; Mostany et al., 2013). Neural structures were imaged using a Zeiss in vivo 2-photon microscope with Zeiss 20x 1.0 NA water-immersion objective lens. High-quality craniotomies had a characteristic bright-field appearance with well-defined vasculature and pale grey matter (Fig. 1A). Under two-photon scanning fluorescent structures were reliably clear and visible with low laser power (<20 mW).

**Figure 1 -.**
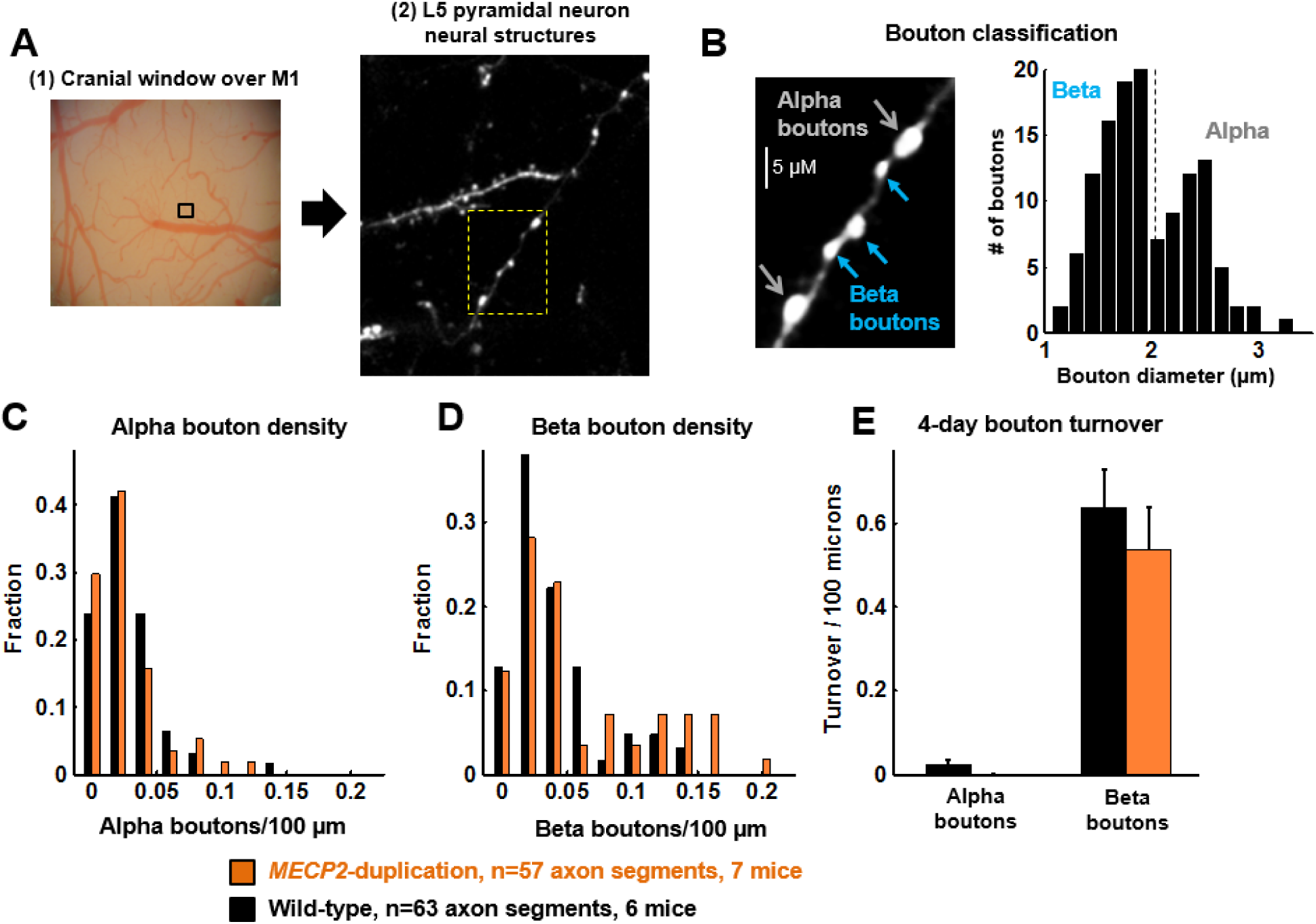
Bouton classification and density of L5 pyramidal neuron axonal projections to layer 1 of mouse primary motor cortex. (**A**): In vivo 2-photon imaging. (1) A cranial window is drilled centered 1.6 mm lateral to the bregma to expose area M1. Correct localization to the forelimb was confirmed post-hoc by electrical microstimulation; see (Ash et al., 2017). (2) GFP-labeled pyramidal neuron processes in layer 1 of motor cortex are imaged. Yellow box shown at high zoom in panel B. (**B**): Bouton classification. <u>*Left:*</u> Varicosities along axons are classified as alpha (>2 μm diameter, blue arrows) or beta (<2 μm diameter, yellow arrows) boutons based on size (see Methods). Extraneous fluorescence structures masked for illustration purposes only. <u>*Right:*</u> Histogram of bouton diameters measured in a subset of axons (n=54 alpha, 74 beta boutons), demonstrating a bimodal distribution. Vertical line depicts the criterion we used to separate alpha from beta boutons. (**C,D**): Histogram of densities of alpha (**C**) and beta (**D**) boutons per axonal segment in *MECP2*-duplication mice (orange,n=57 segments from 7 mice) and WT littermates (black, n=63 segments from 6 mice). (**E**): four-day spontaneous bouton turnover rate, (boutons formed + boutons eliminated) / 2*axon length, for alpha boutons and beta boutons. Alpha boutons were highly stable in this time frame.

Only high quality preparations (low background noise across all time points, <5 pixel i.e. <0.5μm slow motion artifact, <2 pixel i.e. <0.2 μm fast motion artifact, and axons well isolated from other fluorescent structures) were used in the blinded analysis. Pyramidal neuron axons were imaged at high resolution (310x310 to 420x420 μm FOV, 0.1 μm/pixel, 1 μm Z-step size) to adequately capture individual boutons. Laser power was maintained under 20 mW (average ~10 mW) during image stack acquisition.

### Motor training

The Ugo Basile mouse rotarod was used for motor training. At least two hours after imaging sessions, in the late afternoon, mice were placed on the rotarod, and the rotarod gradually accelerated from 5 to 80 rpm over 3 minutes. Single-trial rotarod performance was quantified as the time right before falling or holding on to the dowel rod for two complete rotations without regaining footing. A 7-10 minute rest period occurred between each trial. Four trials were performed per day.

### Analysis of bouton plasticity

<u>*Analysis was performed blind to genotype*</u>. Axons were chosen from the imaging field based on characteristic appearance, including the absence of dendritic spines, minimal branching, and the presence of synaptic boutons, as well as decreased width compared to dendrites. In the thy1-GFP M mouse line (Feng et al., 2000) we employed, the vast majority of GFP-labeled axons in the cerebral cortex arise from L5 pyramidal neurons, though occasional L2/3, L6 pyramidal neurons and thalamocortical neurons may also be labeled (De Paola et al., 2006). Pyramidal neuron axons were targeted based on their thin shafts, high density of small (<1 μm diameter) en-passant boutons, low tortuosity, and rare branching (type A3 axons), allowing them to be clearly distinguished from i) L6 pyramidal neuron axons, which have high branching and a high density of terminaux boutons, and from ii) thalamocortical neurons, which have thicker axons and high branching (De Paola et al., 2006). Given the very sparse labeling of L2/3 neurons in the thy1-GFP M mouse line, we are confident that the great majority of axonal segments we imaged represent L5 pyramidal neuron projections to area L1 from other regions, i.e. chiefly from the premotor, the somatosensory and the contralateral motor cortex (Hooks et al., 2013).

Segments of axon that were clearly visualized in all three time points were selected for analysis (length range 30 – 360 μm, mean 138 μm). The presence of en-passant boutons or terminaux boutons was noted by a blinded investigator, who further classified synaptic boutons as alpha (> ~2 μm or 20 pixel diameter) or beta (<~2 μm or 20 pixel diameter). The threshold used for bouton classification was based on the bimodal distribution of boutons, separable at ~2 μm diameter, present in the analyzed data set (Grillo et al., 2013). The presence of a bouton was determined by a clear increase in axon diameter and increased fluorescence compared to the background axon, as well as the judgment of an experienced investigator (Fig. 1B). Only varicosities that were more than twice as bright as the axonal backbone and extended at least 3 pixels (~0.3 microns) outside the axonal shaft diameter, which corresponds approximately to 2 SDs of the noise blur on either side of the axonal shaft, were counted as boutons. This is similar to (Grillo et al., 2013), which analyzed boutons that were twice as bright as the axon.

Boutons located greater than 50 μm away from the nearest other bouton were excluded from the analysis, so that stretches of bouton-free axon would not bias bouton density calculations. Four to twenty axons were analyzed from 1-3 imaging fields per mouse for 13 mice (6 WT, 7 *MECP2*-duplication mice). Unless the investigator could clearly trace the continuity of axon segments, segments were analyzed as individual units. Though unlikely, the possibility cannot be completely excluded that, on occasion, more than one segment from a single axon were counted. Bouton formation and elimination (Fig. 1C, 2B,C) was calculated as (boutons formed or boutons eliminated) / (total number of boutons observed across imaging sessions), analogous to the measure used in (Grillo et al., 2013). Bouton survival was calculated as the percent of boutons identified in the first imaging time point that are present in subsequent imaging time points. Bouton stabilization was calculated as the percent of newly formed boutons in the second imaging time point, which persisted in the third imaging time point.

**Figure 2 -.**
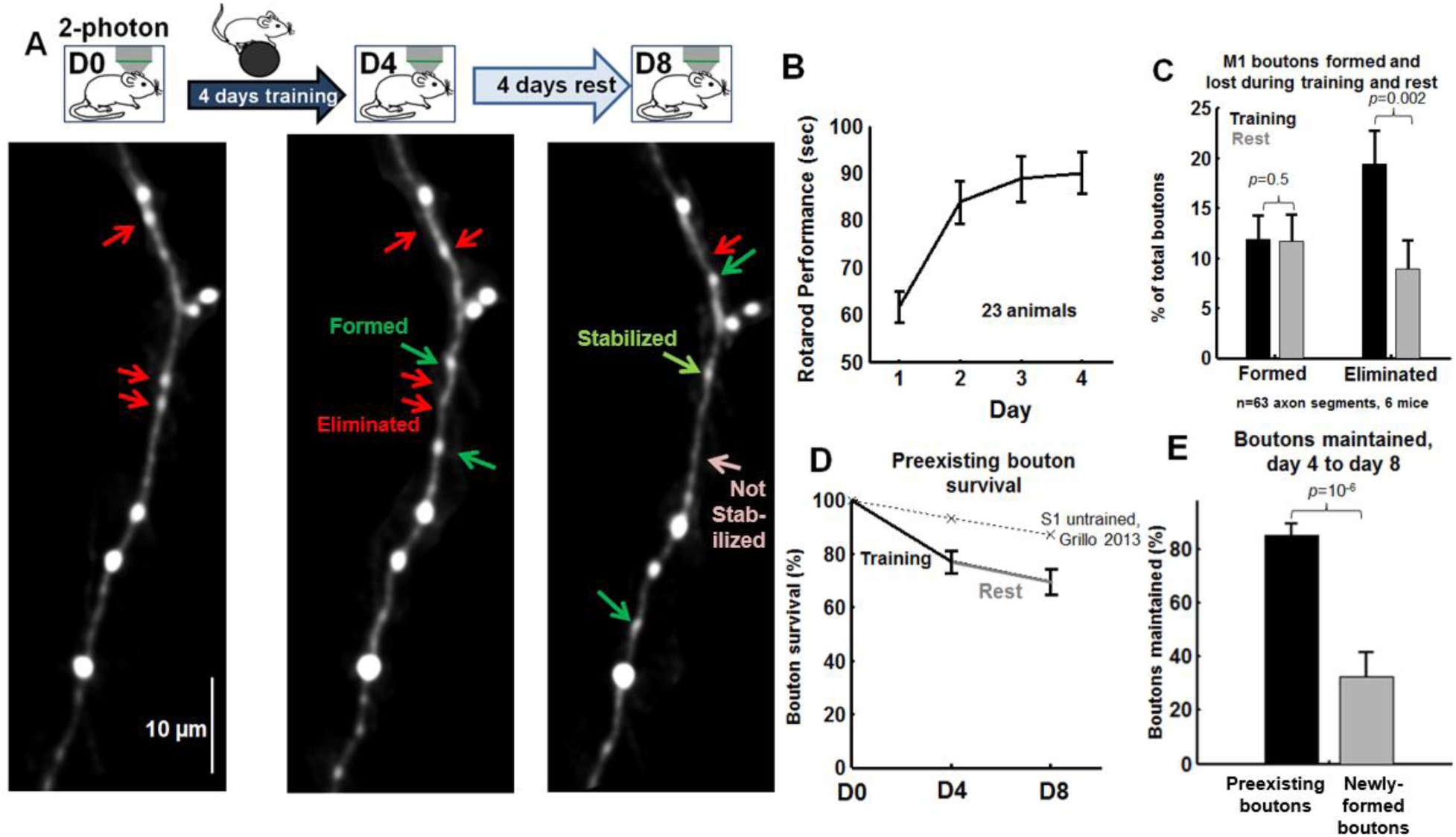
Bouton elimination increases during motor learning in L1 of motor cortex. (**A**): Experimental paradigm and imaging time points. Sample images of axonal segments imaged before (left) and after (middle) 4 days of rotarod training to identify axonal bouton formation (green arrow) and elimination (red arrow) during training. Segments are imaged again following 4 days rest (right) to identify boutons formed, eliminated, and maintained during rest, and learning-associated boutons that are stabilized (light green) or not stabilized (pink). Extraneous fluorescence structures masked and image slightly smoothed for illustration purposes only. (**B**): Median per-day rotarod performance over trials, measured in seconds spent on the accelerating rotarod before falling. Each animal underwent four rotarod training trials per day for 4 days of training (n=23). (**C**): Bouton formation and elimination during training (black) and during rest (grey). Bouton elimination was significantly elevated during training, p=0.001, n=63 segments, Mann-Whitney U test. 314 baseline boutons, 40 formed during training, 42 formed during rest, 64 eliminated during training, 23 eliminated during rest. Data acquired from 6 mice. Statistics performed across axonal segments (p = 0.07 when calculated across animals). (**D**): Pre-existing bouton survival curves across imaging days. Dotted line depicts baseline bouton survival, reproduced from (Grillo et al., 2013). (**E**): The fraction of boutons maintained during the rest period, measured for pre-existing boutons (present on day 0) that were still present on day 4 following training (black) and boutons formed during training (training-associated boutons, grey). p=10^−6^, Mann-Whitney U test.

### Statistics

Except where indicated, the Mann-Whitney U test was used for two-group statistical comparisons, and 2-way ANOVA with Tukey multiple-comparison correction was used for multi-group comparisons.

## RESULTS [1460 words – how many allowed?]

The Tg1 mouse model for *MECP2* duplication syndrome (FVB background) was crossed to the thy1-GFP-M mouse line (C57 background) to generate F1 hybrid males for experiments. A cranial window was placed over motor cortex (1.6 mm lateral to bregma) at 12-14 weeks of age, and at least 2 weeks following the surgery the mouse was placed under the 2-photon microscope to image GFP-labeled axons in layer 1 of area M1 (Fig. 1A; see methods).

L5 pyramidal neuron axons are typically visualized as a thin string of fluorescence interspersed with fluorescent expansions or varicosities (en passant boutons) and rare spine-like terminaux boutons. They are readily differentiated morphologically from L6 neuron axons and thalamocortical axons (De Paola et al., 2006), which, in any case, are rarely fluorescent in these animals. The thy1-GFP M line primarily labels L5 pyramidal neurons in neocortex, and therefore the majority of axonal arbors we imaged are expected to arise from L5 of the somatosensory cortex, the premotor cortex, or the contralateral motor cortex, all of which project to L1 of area M1 (Colechio and Alloway, 2009; Mao et al., 2011; Hooks et al., 2013). Area M1 L5 neurons rarely send projections locally to layer 1 (Cho et al., 2004).

First, we report on axonal bouton structure and plasticity analyzed in littermate controls with normal MECP2 expression. Axonal boutons were identified as periodic thickenings or extensions along the axon, at least twice as bright as and extending at least 0.3 μm from the axonal backbone (Grillo et al., 2013) (this corresponds to approximately 2 SDs of the noise blur on either side of the axonal shaft). We observed a bimodal distribution of bouton sizes, the two modes separated at approximately 2μm diameter (Fig. 1B, right panel). These large (alpha) and small (beta) boutons were analyzed separately. The density of alpha boutons was 2.7±0.3 boutons/100μm (mean±SEM, n=63 axonal segments), and the density of beta boutons was 4.0±0.4 boutons/100μm (Fig. 1C), similar to a previous study (see Methods, Grillo et al., 2013). As expected given their large size (Grillo et al., 2013), alpha boutons were much more stable than beta boutons (Fig. 1D). Across 4 days of rest the 4-day turnover rate (TOR = (gain rate+loss rate) /2) of alpha boutons was 0.5±0.25%, while the TOR of beta boutons was 23±4%. These results are comparable to a previous study in somatosensory cortex, which found 0.1±0.06% 4-day turnover for large boutons and 30±3% 4-day turnover for small boutons (see Fig. 4E,F in Grillo et al., 2013). Since alpha boutons were stable over time, hardly changing over the time course of the experiment, we restricted further analysis of structural plasticity to beta boutons.

**Figure 4 -.**
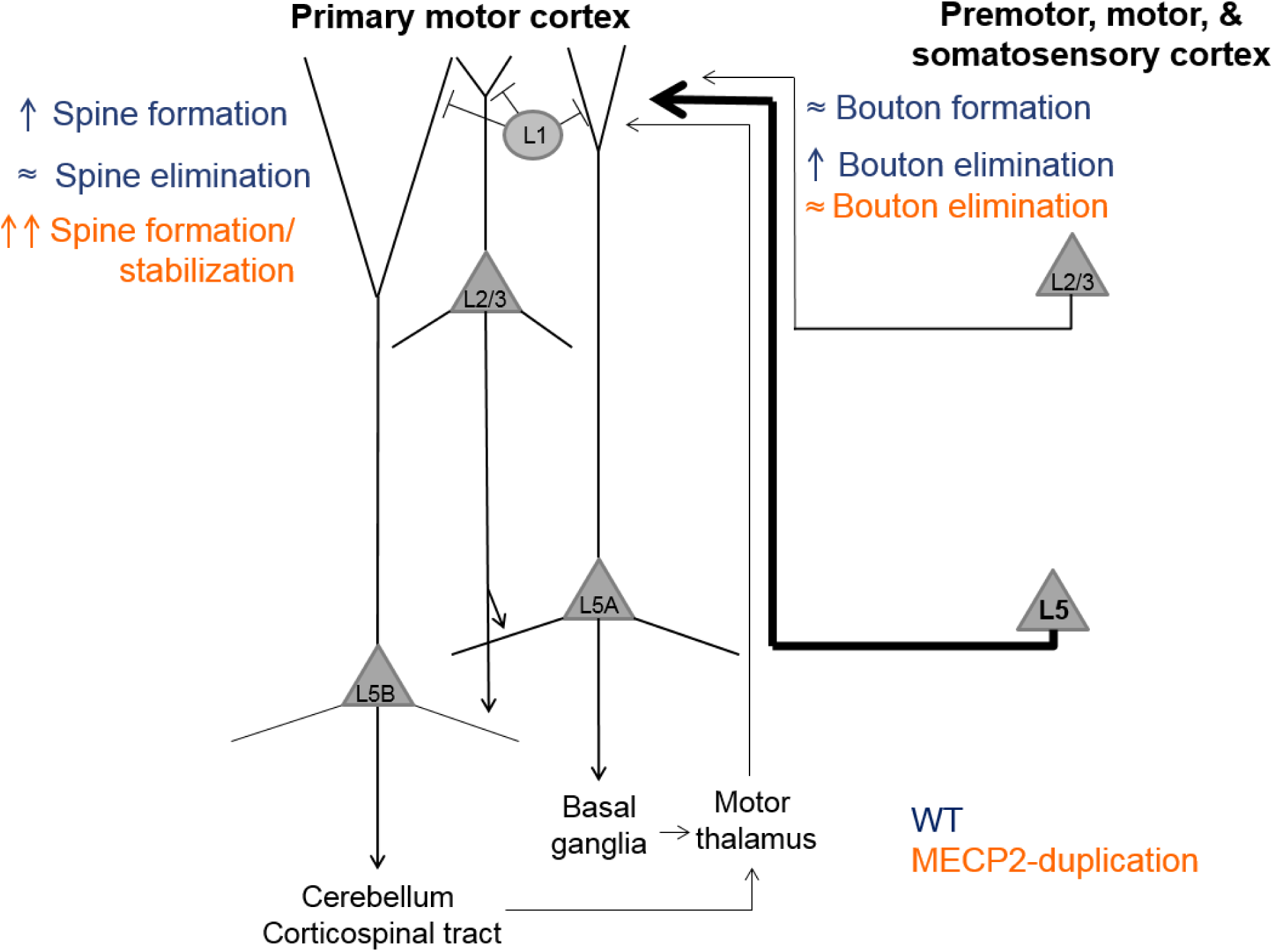
Sketch of structural plasticity phenotypes in dendrites and axonal projections in area M1 of *MECP2*-duplication and WT mice. A highly simplified diagram of the layer 1 motor cortex circuit, including major local connections, inputs, and outputs. The imaged input projection is shown on the right in bold and represents axonal projections to layer 1 from L5 pyramidal neurons in somatosensory, premotor, and contralateral motor cortex. In WT mice (navy blue), spine formation increases in L5B neuron apical dendrites during motor training, while bouton elimination increases in L5 axonal projections. In *MECP2*-duplication mice (orange), spine formation/stabilization increases even more than WT during training, while bouton elimination is unchanged. See text for detail.

The experimental design is diagrammed in Fig. 2A. L5 pyramidal neuron axonal projections to layer 1 (L1) of area M1 were initially imaged to identify baseline boutons. Then mice underwent four days of training on the accelerating rotarod task. Axons were re-imaged to quantify learning-associated bouton turnover. Mice rested in the home cage for four days, and axons were imaged again to observe bouton turnover during rest. WT mice performed progressively better on the rotarod across 4 days of training as reported before (Fig. 2B; see also (Buitrago et al., 2004; Collins et al., 2004)). Training on the rotarod did not significantly alter the formation rate of beta boutons (Fig. 2C; training: 12±2% of total boutons across time points, rest: 12±3% of total boutons, p= 0.5, Mann-Whitney U test n=63 axon segments from 6 mice). The measured formation rate was comparable to the spontaneous 4-day bouton formation rate previously observed in L5 pyramidal neuron axons in somatosensory cortex (8±1%, Fig. S4C in Grillo et al., 2013). Interestingly, rotarod training led to a dramatic increase in bouton elimination compared to rest: 19±3% of total boutons were lost after 4 days of training compared to 9±3% of total boutons lost after 4 days of rest, p = 0.002, Mann-Whitney U test across axonal segments; p = 0.07, paired t test across animals). The M1 spontaneous elimination rate we observed was also comparable to prior published results in area S1 (8.0±0.2%, Grillo et al., 2013, Fig. S4D). Overall, in control animals, motor learning induces a doubling of bouton elimination in M1 without a concomitant change in the rate of bouton formation.

Plotting the survival fraction of pre-existing (“baseline”) boutons revealed that putative L5 pyramidal axons projecting to L1 of area M1 maintained 77±4% of their baseline boutons (boutons present pre-training, on day 0) through 4 days of training (Fig. 2D). This value is significantly lower than prior estimates of spontaneous 4-day survival fraction of L5 pyramidal neuron axonal boutons (~90% of baseline boutons, dotted line in Fig. 2D, see Fig. 7B of De Paola et al., 2006, Fig. 3C of Grillo et al., 2013, Fig. 5 of Majewska et al., 2006). Note that elimination rates (Fig. 1C) and survival curves (Fig. 1D) do not sum exactly to 100% because elimination rate was calculated as a fraction of the total number of boutons observed across all time points to avoid outlier turnover rates in axons which had very few baseline boutons, following Grillo et al., 2013 (see Methods).

**Figure 3 -.**
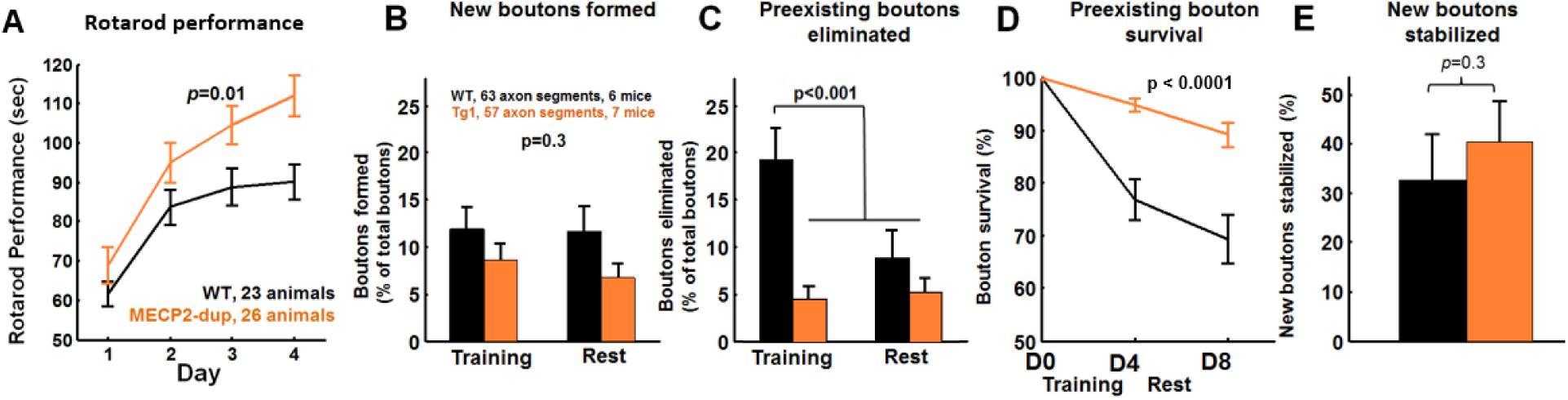
Increased stability of axonal boutons during training in *MECP2*-duplication mice. (**A**): Median per-day rotarod performance averaged across animals, measured by the time (seconds) spent on the accelerating rotarod before falling. Four trials were performed per day across 4 days of training, in *MECP2*-duplication (orange, n=26) and WT (black, n=23) animals (Collins et al., 2004). Significance was assessed by repeated-measures ANOVA. This data is reproduced from (Ash et al., 2017) and illustrates that training was effective. (**B**): Bouton formation during training (training-associated boutons) and during rest in *MECP2*-duplication mice and WT littermates, p=0.3, 2-way ANOVA. (**C**): Pre-existing bouton elimination during training and during rest in each genotype, p< 0.001, bouton elimination during rest in WT vs all other conditions, 2-way ANOVA with Tukey test for multiple comparisons. Effect of genotype: F=12.9, p=0.0004; effect of training vs. rest: F=5.3, p = 0.02; genotype x training interaction: F=3.71, p = 0.055. Linear mixed-effects model ANOVA results: Effect of genotype: t=-2.4, p=0.015; effect of training vs. rest: t=-2.6, p = 0.009; genotype x training interaction: t=1.9, p = 0.053. (**D**): Pre-existing bouton survival curves across imaging, p<0.0001 effect of genotype, 2-way ANOVA. Effect of genotype: F=26.7, p<0.0001; effect of training vs. rest: F=3.25, p = 0.07; genotype x training interaction: F=0.06, p = 0.8. (**E**): Training-associated bouton stabilization rate – the number of boutons formed during training and still present after 4 days of post-training rest is not significantly different across genotypes. Data are plotted as percentage of boutons formed during training. Mann-Whitney U test.

We also compared the survival rate of newly formed training-related boutons with that of pre-existing boutons. In the four days of rest following training, 85±4% of baseline preexisting boutons (boutons present on day zero that were also present on post-training day 4) were maintained, while newly formed boutons were maintained at a much lower rate of 32±9% (Fig. 2E, p=10^−6^), consistent with the reported stabilization rate of spontaneously formed boutons in somatosensory cortex (newly formed: 35±5% of all boutons over 4 days, Grillo et al., 2013).

We then assessed learning-associated axonal bouton turnover in *MECP2*-duplication mice. *MECP2*-duplication mice performed significantly better on the rotarod than control littermates (Fig. 3A) as previously described (Collins et al., 2004; Ash et al., 2017). The average length of analyzed axonal segments was not significantly different between mutants and WT littermates (WT: 142±73 μm, *MECP2*-duplication: 133±73 μm, mean±SD). The density of alpha boutons (Fig. 1C) and beta boutons (Fig. 1D) was also similar between the genotypes (<u>*alpha boutons*</u>, control: 2.7±0.3 boutons/100μm, *MECP2*-duplication: 2.4±0.3 boutons/100μm, p=0.4; *<u>beta boutons</u>*, control: 4±0.4 boutons/100μm, *MECP2*-duplication: 5.8±0.7 boutons/100μm. p=0.2, Mann-Whitney U test). Similar to WT, alpha boutons were highly stable compared to beta boutons in *MECP2*-duplication mice (data not shown). The rate of beta bouton formation was not significantly different between *MECP2*-duplication mice and WT controls, neither during the training (Fig. 3B, control: 12±2% of total boutons, n=63 axon segments from 6 mice; *MECP2*-duplication: 9±3% of total boutons, n=57 axon segments from 7 mice) nor during the rest phase (control: 12±3 % of total boutons, *MECP2*-duplication: 7±1 % of total boutons, effect of genotype: F=2.8, p =0.09; effect of training vs. rest: F=0.78, p=0.4; genotype x training interaction: F=0.6, p=0.4). Interestingly, the increased bouton elimination rate during training observed in WT mice (training: 19±3%, rest: 9±3% of total boutons) did not occur in *MECP2*-duplication mice (Fig. 3C, training: 5±1% of total boutons; rest: 5±1% of total boutons). Significantly fewer boutons were eliminated during training in *MECP2*-duplication mice compared to littermate controls (p < 0.001, bouton elimination during training in WT vs. all other groups, 2-way ANOVA with Tukey-test for multiple comparisons. Effect of genotype: F=12.9, p=0.0004; effect of training vs. rest: F=5.3, p = 0.02; genotype x training interaction: F=3.71, p = 0.055). A linear mixed-effects model ANOVA, with genotype and imaging time point implemented as fixed effects, and mouse implemented as a random effect generated similarly significant results (Effect of genotype: t=-2.4, p=0.015; effect of training vs. rest: t=-2.6, p = 0.009; genotype x training interaction: t=1.9, p = 0.053).

Plotting the survival fraction of boutons revealed that baseline boutons were significantly more stable in *MECP2*-duplication mice vs. littermate controls (Fig. 3E, p < 0.0001, 2-way ANOVA. Effect of genotype: F=26.7, p<0.0001; effect of training vs. rest: F=3.25, p = 0.07; genotype x training interaction: F=0.06, p = 0.8). *MECP2*-duplication axons maintained 95±1% of their boutons after 4 days of training, while control littermate axons maintained only 77±4%. Subsequently, in the four days of rest following training, bouton loss was comparable between genotypes. *MECP2*-duplication axons lost a further 6±1% of baseline boutons to reach 89±2% bouton survival on day eight, while littermate controls lost a further 8±2% to end at 69±4%. Learning-associated bouton stabilization rate, defined as the fraction of boutons formed during the four days of training that were still present after four further days of rest, was not significantly altered in *MECP2*-duplication mice (40±8%) comped to controls (32±9%, Fig. 3F, p = 0.3). Again, note that elimination rates (Fig. 3C) and survival curves (Fig. 3D) do not sum to 100%, as explained above, but note that the measured differences remain significant if the elimination rate is calculated as a fraction of baseline boutons instead of as a fraction of total boutons across time points.

Bouton formation, elimination, and stabilization rates did not correlate well with rotarod performance in individual animals for either genotype (p > 0.05, t test on linear regression, all comparisons, data not shown), suggesting that other factors are potentially more important for the behavioral manifestations of motor learning.

## DISCUSSION

The stability and plasticity of synaptic connections is a tightly regulated process that unfolds throughout life. A pathological imbalance between stability and plasticity could lead to the altered patterns of learning and forgetting observed in autism mouse models (Collins et al., 2004; Rothwell et al., 2014) and in autistic patients (Treffert, 2014). In prior work (Ash et al., 2017) an abnormal increase in learning-associated dendritic spine stability was found after motor training in the apical tuft of area M1 corticospinal neurons in the Tg1 mouse model of *MECP2* duplication syndrome. Here we investigated how axonal boutons in the L5 pyramidal neuron projection to L1 of primary motor cortex turn over during motor training in these animals. First, we find in WT mice that: **1)** bouton formation rate is unaffected by motor learning (Fig.2 C), and **2)** bouton elimination rate doubles from ~10% to ~20% during motor learning (Fig. 2C,D). In contrast, we find that the increase in learning-associated bouton elimination observed in littermate controls does not occur in *MECP2*-duplication mice (Fig. 3C), which exhibit increased bouton stability during training (Fig. 3D). Bouton formation rate during motor learning was similar between *MECP2*-duplication animals and littermate controls (Fig. 3B), and was not significantly different from the rate of bouton formation observed at rest in either genotype. A similar fraction of learning-associated boutons was stabilized in both genotypes (Fig. 3E).

### Bouton formation and elimination with motor learning in controls

Our spontaneous 4-day bouton turnover results are in agreement with a previous study of axonal bouton formation and elimination in L5 pyramidal neuron axons projecting to layer 1 of somatosensory cortex (Grillo et al., 2013), suggesting that baseline axonal bouton turnover in L1 is similar in sensory and motor areas. Here, we found that, in normal animals, the rate of axonal bouton elimination increases markedly during motor training in L5 pyramidal neuron projections to L1 of area M1, without a concomitant increase in the rate of bouton formation (Fig. 2C).

At face value this suggests that training leads to a weakening of L5 pyramidal inputs to layer 1 of the primary motor cortex, at least as evidenced by structural analysis. L5 Axonal projections to L1 have several potential synaptic partners, including apical dendritic arbors of L5B corticospinal pyramidal neurons, L5A corticostriatal/corticocallosal neurons, L2/3 pyramidal neurons, and L1 interneuron dendrites (Fig. 4). Since L1 interneurons are sparse, most of the postsynaptic partners of the axonal boutons we studied are likely formed with one or more of the aforementioned classes of pyramidal neurons. The increased elimination of pre-synaptic axonal boutons would then lead us to expect a corresponding loss in their post-synaptic partners, i.e. of dendritic spines located in the apical dendritic tufts of the target neurons. However, we and others have previously observed an increase in the formation rate of dendritic spines during motor learning in the apical tuft terminal dendrites of L5 neurons in layer 1 of area M1 (Xu et al., 2009; Yang et al., 2009; Liston et al., 2013; Ash et al., 2017) without an increase in spine elimination during training. This dissociation between L5 neuron dendritic spine formation and axonal bouton elimination in layer 1 of area M1 during motor learning suggests that at least a subset of the axonal projections we imaged are synapsing on dendritic processes not previously analyzed. Indeed, there is evidence that projections to L1 of M1 from different brain areas preferentially target different cell types (Hooks et al., 2013). Ergo it is also possible that the pre-synaptic partners of the L5 apical tuft dendritic spines studied previously during motor learning (Xu et al., 2009; Yang et al., 2009) may arise from thalamocortical, L2/3, or L6 projections which we did not study here. Overall, these results raise the interesting possibility that different pathways projecting to L1 of mouse area M1 may have different signatures of structural plasticity during motor learning.

We note the alternative possibility that the axonal boutons (or varicosities) whose elimination rate increases with learning are ones that have not yet formed a synapse, and therefore do not have post-synaptic partners. It has been estimated that ~10-20% of axonal varicosities (defined as a swelling of the axon exceeding the shaft diameter by more than 50%) may be non-synaptic (Shepherd and Harris, 1998; White et al., 2004; Bourne et al., 2013), although this has not been assessed in pyramidal neuron projections to layer 1 to our knowledge.

This still leaves us with the puzzle of why we do not observe an increase in the rate of bouton formation with learning, to serve as pre-synaptic partners to the increased number of dendritic spines that form in apical dendritic tufts of L5 pyramidal neurons (Fig. 4). One possibility mentioned above is that we have not examined the correct pre-synaptic axons, particularly as we did not study thalamo-cortical, L2/3 or L6 pyramidal projections. Another possibility is that, rather than connecting with a new axonal bouton, newly formed spines rather form a second synapse onto large pre-existing boutons already harboring a synapse, as has been shown in vivo in the somatosensory cortex and ex vivo in the hippocampus (~70% of newly formed spines synapse with a multi-synapse bouton, compared to 20-30% of preexisting spines (Knott et al., 2006; Nagerl et al., 2007); see also (Woolley et al., 1996; Toni et al., 1999; Geinisman et al., 2001; Yankova et al., 2001; Federmeier et al., 2002; Nicholson and Geinisman, 2009; Lee et al., 2013). Dendritic spines formed during learning may largely synapse on already existing, large, pre-synaptic boutons (alpha boutons in our study) where they compete with the previously present connections. Over time, some of these connections withdraw, re-establishing a new equilibrium that favors the new skill learning. Presumably, in the days-to-weeks following learning, bouton formation modestly increases and/or bouton elimination decreases to bring bouton densities back to baseline levels.

#### Increased bouton stability in *MECP2*-duplication mice

We found that the learning-associated increase in bouton elimination rate occurring in WT mice is abolished in *MECP2*-duplication mice. This increased synaptic stability in L5 pyramidal neuron axonal projections could reflect a general increase in synaptic stability during motor learning across multiple circuits, or it may be a manifestation of increased stability along specific subcircuits projecting to L1 of area M1 (Fig. 4). Increased bouton stability in *MECP2*-duplication M1 could also be due in part to more robust capture and stabilization of pre-existing boutons by newly formed learning-associated spines, boutons that would have otherwise been eliminated due to loss of their prior post-synaptic targets during the training period (Knott et al., 2006; Nagerl et al., 2007). Quantification of multi-synapse bouton density with and without training in mutants could address these possibilities. As in the WT, again it is interesting to note that the learning-associated increase in spine formation in mutants is not associated with a concomitant increase in bouton formation (Fig. 4). This provides further basis for the idea that different microcircuit projections to M1 may manifest different patterns of structural synaptic plasticity during learning.

It is interesting to speculate that the learning-associated bouton elimination that occurs in littermate controls is a natural end result of strong long-term depression (Becker et al., 2008; Wiegert and Oertner, 2013). In this case, the lack of bouton elimination in mutants may connote a disruption in processes regulating LTD. Taken along with the fact that abnormal LTD is observed in many other autism models (D’Antoni et al., 2014), it will be interesting to experimentally test if LTD is indeed altered in motor cortex of *MECP2*-duplication mice, and to see if decreased LTD underlies the mutant’s increased learning-associated bouton stability.

The behavioral implications of increased L1 axonal bouton stabilization remain a matter of speculation. The rate of pre-existing bouton formation and elimination did not correlate with behavioral performance across individual animals in either genotype. This suggests that although bouton elimination is a robust structural phenotype resulting from motor training, its link to behavioral performance is at best weak. It is certainly weaker than dendritic spine formation and stabilization in apical dendrites of L5 pyramidal neurons, which correlates well with behavior (Yang et al., 2009; Ash et al., 2017). It is possible that our study is not adequately powered to detect a weak correlation between bouton turnover and rotarod performance.

#### Potential Limitations

It is important to note a number of limitations with the study. First of all, our quantification of presynaptic terminals depends entirely on morphological measures. Although we used conservative criteria similar to that which in prior experimenters’ hands has been shown to reliably detect synapse-forming puncta (De Paola et al., 2006), it is still possible that a fraction (~10%) of the counted varicosities are non-synaptic (Shepherd and Harris, 1998; White et al., 2004; Bourne et al., 2013).

Second, the rest phase occurred following training, so it is possible that some of the corresponding bouton turnover may reflect enduring consolidation processes that persist beyond training rather than a true rest phase. Having said that, the measured spontaneous axonal bouton formation and elimination is in very close agreement to previous studies (Grillo et al., 2013), suggesting that the measurements reflect baseline turnover.

Third, we cannot precisely determine the origin of the axonal afferents imaged in our study (Fig. 4). Some of the heterogeneity in plasticity observed across imaged axons could be due to projection-specific differences. For example, it would be interesting to speculate that coarse sensorimotor training induced by the rotarod may drive greater bouton remodeling in somatosensory cortical inputs to area M1, while fine motor training requiring higher-order motor planning, such as the seed-grabbing task used by (Xu et al., 2009), may induce greater remodeling in premotor cortical inputs.

Fourth, the postsynaptic partners of the imaged axons are unknown. The precise connectivity of inputs to M1, with S1 pyramidal neuron axons preferentially synapsing on L2/3 and L5A neurons and premotor cortex pyramidal neuron axons preferentially synapsing on L5B neurons (Mao et al., 2011; Hooks et al., 2013), enables a rich potential repertoire of synaptic reorganization during training. New methods targeting fluorescent proteins to specific input areas, as well as combinatorial techniques labeling pre-and postsynaptic partners (Kim et al., 2011; Druckmann et al., 2014), will enable scientists to tackle this question in the future.

### Conclusions and implications

In conclusion, we report here that L5 pyramidal neuron axonal projections to layer 1 of WT mouse motor cortex exhibit a selective escalation in bouton elimination during motor learning, a plasticity process that is disrupted in the *MECP2*-duplication syndrome mouse model of autism. These data constrain models of motor cortex plasticity underlying learning and underscore the possibility that different synaptic pathways within the cortical circuit may manifest different patterns of structural synaptic plasticity during learning. Future work studying plasticity along different synaptic pathways that link various areas along the motor circuit will shed further light on these issues.

Our results provide further evidence for an altered balance between stability and plasticity of synaptic connections in favor of stability in the *MECP2* duplication syndrome mouse model (Ash et al., 2017). This bias favors enhanced motor learning on the rotarod and may play a role in other types of learning, such as fear conditioning or social learning. More generally, an abnormal bias toward synaptic stability in relevant circuits could potentially play a role in explaining the combination of savant-like phenotypes and behavioral rigidity seen at times in autism.

## Acknowledgments

R.T.A. received support from the Autism Speaks Weatherstone Fellowship and the BCM Medical Scientist Training Program. This work was supported by grants from the Simons Foundation and March of Dimes to S.M.S., the Howard Hughes Medical Institute and NINDS HD053862 to H.Y.Z., and the Baylor Intellectual and Developmental Disabilities Research Center (P30HD024064) Mouse Neurobehavioral Core. We are grateful to S. Torsky, B. Suter, J. Patterson, S. Shen, and D. Yu for technical and theoretical advice on experiments and comments on the manuscript.

## REFERENCES

Antar LN, Li C, Zhang H, Carroll RC, Bassell GJ (2006) Local functions for FMRP in axon growth cone motility and activity-dependent regulation of filopodia and spine synapses. Mol Cell Neurosci 32:37–48.

Antonova I, Arancio O, Trillat a C, Wang HG, Zablow L, Udo H, Kandel ER, Hawkins RD (2001) Rapid increase in clusters of presynaptic proteins at onset of long-lasting potentiation. Science 294:1547–1550.

Ash RT, Buffington SA, Park J, Costa-Mattioli M, Zoghbi HY, Smirnakis SM (2017) Excessive ERK-dependent synaptic clustering with enhanced motor learning in the MECP2 duplication syndrome mouse model of autism. bioRxiv.

Becker N, Wierenga CJ, Fonseca R, Bonhoeffer T, Nägerl UV (2008) LTD Induction Causes Morphological Changes of Presynaptic Boutons and Reduces Their Contacts with Spines. Neuron 60:590–597.

Belichenko P V., Wright EE, Belichenko NP, Masliah E, Li HH, Mobley WC, Francke U (2009) Widespread changes in dendritic and axonal morphology in Mecp2-mutant mouse models of Rett syndrome: Evidence for disruption of neuronal networks. J Comp Neurol 514:240–258.

Blundell J, Kaeser PS, Südhof TC, Powell CM (2010) RIM1 and Interacting Proteins Involved in Presynaptic Plasticity Mediate Prepulse Inhibition and Additional Behaviors Linked to Schizophrenia. J Neurosci 30:5326–5333 Available at: http://www.jneurosci.org/cgi/doi/10.1523/JNEUROSCI.0328-10.2010%5Cnpapers3://publication/doi/10.1523/JNEUROSCI.0328-10.2010.

Bourne JN, Chirillo MA, Harris KM (2013) Presynaptic ultrastructural plasticity along CA3???CA1 axons during long-term potentiation in mature hippocampus. J Comp Neurol 521:3898–3912.

Buitrago MM, Schulz JB, Dichgans J, Luft AR (2004) Short and long-term motor skill learning in an accelerated rotarod training paradigm. Neurobiol Learn Mem 81:211–216.

Carrillo J, Cheng S-Y, Ko KW, Jones T a, Nishiyama H (2013) The long-term structural plasticity of cerebellar parallel fiber axons and its modulation by motor learning. J Neurosci 33:8301–8307 Available at: http://www.pubmedcentral.nih.gov/articlerender.fcgi?artid=3680104&tool=pmcentrez&rendertype=abstract.

Chen J, Yu S, Fu Y, Li X (2014) Synaptic proteins and receptors defects in autism spectrum disorders. Front Cell Neurosci 8:276 Available at: http://www.pubmedcentral.nih.gov/articlerender.fcgi?artid=4161164&tool=pmcentrez&rendertype=abstract.

Chklovskii DB, Mel BW, Svoboda K (2004) Cortical rewiring and information storage. Nature 431:782–788.

Cho RH, Segawa S, Okamoto K, Mizuno A, Kaneko T (2004) Intracellularly labeled pyramidal neurons in the cortical areas projecting to the spinal cord: II. Intra-and juxta-columnar projection of pyramidal neurons to corticospinal neurons. Neurosci Res 50:395–410.

Colechio EM, Alloway KD (2009) Differential topography of the bilateral cortical projections to the whisker and forepaw regions in rat motor cortex. Brain Struct Funct 213:423–439.

Collingridge GL, Peineau S, Howland JG, Wang YT (2010) Long-term depression in the CNS. Nat Rev Neurosci 11:459–473.

Collins AL, Levenson JM, Vilaythong AP, Richman R, Armstrong DL, Noebels JL, Sweatt D, J, Zoghbi HY (2004) Mild overexpression of MeCP2 causes a progressive neurological disorder in mice. Hum Mol Genet 13:2679–2689.

D’Antoni S, Spatuzza M, Bonaccorso CM, Musumeci SA, Ciranna L, Nicoletti F, Huber KM, Catania MV (2014) Dysregulation of group-I metabotropic glutamate (mGlu) receptor mediated signalling in disorders associated with Intellectual Disability and Autism. Neurosci Biobehav Rev 46:228–241.

De Paola V, Holtmaat A, Knott G, Song S, Wilbrecht L, Caroni P, Svoboda K (2006) Cell type-specific structural plasticity of axonal branches and boutons in the adult neocortex. Neuron 49:861–875.

Degano AL, Pasterkamp RJ, Ronnett G V. (2009) MeCP2 deficiency disrupts axonal guidance, fasciculation, and targeting by altering Semaphorin 3F function. Mol Cell Neurosci 42:243–254.

Deng PY, Rotman Z, Blundon JA, Cho Y, Cui J, Cavalli V, Zakharenko SS, Klyachko VA (2013) FMRP Regulates Neurotransmitter Release and Synaptic Information Transmission by Modulating Action Potential Duration via BK Channels. Neuron 77:696–711.

Druckmann S, Feng L, Lee B, Yook C, Zhao T, Magee JC, Kim J (2014) Structured Synaptic Connectivity between Hippocampal Regions. Neuron 81:629–640.

Federmeier KD, Kleim JA, Greenough WT (2002) Learning-induced multiple synapse formation in rat cerebellar cortex. Neurosci Lett 332:180–184.

Feng G, Mellor RH, Bernstein M, Keller-Peck C, Nguyen QT, Wallace M, Nerbonne JM, Lichtman JW, Sanes JR (2000) Imaging neuronal subsets in transgenic mice expressing multiple spectral variants of GFP. Neuron 28:41–51 Available at: http://www.ncbi.nlm.nih.gov/pubmed/11086982.

Garcia-Junco-Clemente P, Golshani P (2014) PTEN: A master regulator of neuronal structure, function, and plasticity. Commun Integr Biol 7.

Geinisman Y, Berry RW, Disterhoft JF, Power JM, Van der Zee E a (2001) Associative learning elicits the formation of multiple-synapse boutons. J Neurosci 21:5568–5573.

Grillo FW, Song S, Teles-Grilo Ruivo LM, Huang L, Gao GG, Knott GW, Maco B, Ferretti V, Thompson D, Little GE, De Paola V (2013) Increased axonal bouton dynamics in the aging mouse cortex. Proc Natl Acad Sci 110:1–10 Available at: http://www.pnas.org/cgi/doi/10.1073/pnas.1218731110%5Cnhttp://www.pubmedcentral.nih.gov/articlerender.fcgi?artid=3631669&tool=pmcentrez&rendertype=abstract.

Holtmaat A, Bonhoeffer T, Chow DK, Chuckowree J, De Paola V, Hofer SB, Hübener M, Keck T, Knott G, Lee W-CA, Mostany R, Mrsic-Flogel TD, Nedivi E, Portera-Cailliau C, Svoboda K, Trachtenberg JT, Wilbrecht L (2009) Long-term, high-resolution imaging in the mouse neocortex through a chronic cranial window. Nat Protoc 4:1128–1144 Available at: http://www.nature.com/doifinder/10.1038/nprot.2009.89 [Accessed February 16, 2017].

Hooks BM, Mao T, Gutnisky DA, Yamawaki N, Svoboda K, Shepherd GMG (2013) Organization of Cortical and Thalamic Input to Pyramidal Neurons in Mouse Motor Cortex. J Neurosci 33:748–760 Available at: http://www.jneurosci.org/content/33/2/748.long%5Cnhttp://www.jneurosci.org/cgi/doi/10.1523/JNEUROSCI.4338-12.2013.

Johnson CM, Peckler H, Tai L-H, Wilbrecht L (2016) Rule learning enhances structural plasticity of long-range axons in frontal cortex. Nat Commun 7:10785 Available at: http://www.nature.com/ncomms/2016/160307/ncomms10785/full/ncomms10785.html.

Kim J, Zhao T, Petralia RS, Yu Y, Peng H, Myers E, Magee JC (2011) mGRASP enables mapping mammalian synaptic connectivity with light microscopy. Nat Methods 9:96–102 Available at: http://dx.doi.org/10.1038/nmeth.1784.

Knott GW, Holtmaat A, Wilbrecht L, Welker E, Svoboda K (2006) Spine growth precedes synapse formation in the adult neocortex in vivo. Nat Neurosci 9:1117–1124 Available at: http://www.ncbi.nlm.nih.gov/pubmed/16892056 [Accessed February 16, 2017].

Lee KJ, Park IS, Kim H, Greenough WT, Pak DTS, Rhyu IJ (2013) Motor Skill Training Induces Coordinated Strengthening and Weakening between Neighboring Synapses. J Neurosci 33:9794–9799 Available at: http://www.jneurosci.org/content/33/23/9794%5Cnhttp://libsta28.lib.cam.ac.uk:2356/content/33/23/9794%5Cnhttp://www.jneurosci.org/content/33/23/9794.full.pdf%5Cnhttp://www.ncbi.nlm.nih.gov/pubmed/23739975.

Liston C, Cichon JM, Jeanneteau F, Jia Z, Chao M V, Gan W-B (2013) Circadian glucocorticoid oscillations promote learning-dependent synapse formation and maintenance. Nat Neurosci 16:698–705 Available at: http://dx.doi.org/10.1038/nn.3387.

Majewska AK, Newton JR, Sur M (2006) Remodeling of Synaptic Structure in Sensory Cortical Areas In Vivo. J Neurosci 26.

Mao T, Kusefoglu D, Hooks BM, Huber D, Petreanu L, Svoboda K (2011) Long-Range Neuronal Circuits Underlying the Interaction between Sensory and Motor Cortex. Neuron 72:111–123.

Mostany R, Anstey JE, Crump KL, Maco B, Knott G, Portera-Cailliau C (2013) Altered Synaptic Dynamics during Normal Brain Aging. J Neurosci 33:4094–4104 Available at: http://www.jneurosci.org/cgi/doi/10.1523/JNEUROSCI.4825-12.2013.

Mostany R, Portera-Cailliau C (2008) A Craniotomy Surgery Procedure for Chronic Brain Imaging. J Vis Exp:e680–e680 Available at: http://www.jove.com/index/Details.stp?ID=680 [Accessed September 1, 2016].

Nagerl U V, Kostinger G, Anderson JC, Martin KA, Bonhoeffer T (2007) Protracted synaptogenesis after activity-dependent spinogenesis in hippocampal neurons. J Neurosci 27:8149–8156 Available at: http://www.ncbi.nlm.nih.gov/pubmed/17652605.

Nicholson DA, Geinisman Y (2009) Axospinous synaptic subtype-specific differences in structure, size, ionotropic receptor expression, and connectivity in apical dendritic regions of rat hippocampal CA1 pyramidal neurons. J Comp Neurol 512:399–418.

Ramocki MB, Tavyev YJ, Peters SU (2010) The MECP2 duplication syndrome. Am J Med Genet Part A 152:1079–1088.

Rothwell PE, Fuccillo M V., Maxeiner S, Hayton SJ, Gokce O, Lim BK, Fowler SC, Malenka RC, Südhof TC (2014) Autism-associated neuroligin-3 mutations commonly impair striatal circuits to boost repetitive behaviors. Cell 158:198–212.

Shepherd GM, Harris KM (1998) Three-dimensional structure and composition of CA3-CA1 axons in rat hippocampal slices: implications for presynaptic connectivity and compartmentalization. J Neurosci 18:8300–8310.

Stettler DD, Yamahachi H, Li W, Denk W, Gilbert CD (2006) Axons and synaptic boutons are highly dynamic in adult visual cortex. Neuron 49:877–887.

Tennant KA, Adkins DL, Donlan NA, Asay AL, Thomas N, Kleim JA, Jones TA (2011) The organization of the forelimb representation of the C57BL/6 mouse motor cortex as defined by intracortical microstimulation and cytoarchitecture. Cereb Cortex 21:865–876.

Toni N, Buchs P a, Nikonenko I, Bron CR, Muller D (1999) LTP promotes formation of multiple spine synapses between a single axon terminal and a dendrite. Nature 402:421–425.

Treffert DA (2014) Savant syndrome: Realities, myths and misconceptions. J Autism Dev Disord 44:564–571.

White EL, Weinfeld E, Lev DL (2004) Quantitative analysis of synaptic distribution along thalamocortical axons in adult mouse barrels. J Comp Neurol 479:56–69.

Wiegert JS, Oertner TG (2013) Long-term depression triggers the selective elimination of weakly integrated synapses. Proc Natl Acad Sci U S A 110:E4510–E4519 Available at: http://www.pnas.org/cgi/doi/10.1073/pnas.1315926110.

Woolley CS, Wenzel HJ, Schwartzkroin PA (1996) Estradiol increases the frequency of multiple synapse boutons in the hippocampal CA1 region of the adult female rat. J Comp Neurol 373:108–117.

Xu T, Yu X, Perlik AJ, Tobin WF, Zweig JA, Tennant K, Jones T, Zuo Y (2009) Rapid formation and selective stabilization of synapses for enduring motor memories. Nature 462:915–919 Available at: http://www.ncbi.nlm.nih.gov/pubmed/19946267.

Yang G, Pan F, Gan WB (2009) Stably maintained dendritic spines are associated with lifelong memories. Nature 462:920–924 Available at: http://www.neuroscience.ubc.ca/CourseMat/Yang_et_al_2009.pdf%5Cnhttp://www.ncbi.nlm.nih.gov/pubmed/19946265.

Yankova M, Hart S a, Woolley CS (2001) Estrogen increases synaptic connectivity between single presynaptic inputs and multiple postsynaptic CA1 pyramidal cells: a serial electron-microscopic study. Proc Natl Acad Sci U S A 98:3525–3530 Available at: http://www.pubmedcentral.nih.gov/articlerender.fcgi?artid=30686&tool=pmcentrez&rendertype=abstract.

